# Unidirectional protein recruitment is essential for viral latency

**DOI:** 10.1101/2021.10.10.463816

**Authors:** Ido Lavi, Supriya Bhattacharya, Ankita Awase, Ola Orgil, Nir Avital, Guy Journo, Vyacheslav Gurevich, Meir Shamay

## Abstract

Kaposi’s sarcoma associated herpesvirus (KSHV, HHV-8) is associated with several human malignancies. During latency the viral genomes reside in the nucleus of infected cells as large non-integrated plasmids, known as episomes. All KSHV infected cells express LANA, and LANA is essential for viral latency. LANA binding to the viral episomes is critical for both replication of the viral genomes during latency, and for tethering the viral episomes to the cell chromosomes during cell division. Directional recruitment of protein complexes are critical for proper function of many nuclear processes. To test for recruitment directionality between LANA and cellular proteins we directed LANA via catalytically inactive Cas9 (dCas9) to a repeat sequence to obtain easily detectable dots. Then, recruitment of nuclear proteins to these dots can be evaluated. We found that LANA recruited its known interactors ORC2 and SIN3A. Interestingly, LANA was unable to recruit MeCP2, but MeCP2 recruited LANA. Both LANA and histone deacetylase 1 (HDAC1) interact with the transcriptional-repression domain (TRD) of MeCP2. Similar to LANA, HDAC1 was unable to recruit MeCP2. While heterochromatin protein 1 (HP1) which interacts with the N-terminal of MeCP2, was able to recruit MeCP2. Forcing MeCP2 dimerization via the tandem epitopes of SunTag, allows LANA to recruit MeCP2 in infected cells. We propose that available interacting domains force this recruitment directionality. Knock-down of MeCP2 by shRNA, as well as cells derived from Rett syndrome and express a mutant MeCP2 (T158M), dramatically reduced the ability of LANA to support viral latency. Therefore, this unidirectional recruitment of LANA by MeCP2 identified MeCP2 as a critical factor for KSHV viral maintenance.

**Significance Statement:** Using a CRISPR/Cas9 recruitment assay, we show that some interacting proteins have a unidirectional recruitment property, where only one of the proteins can recruit its partner. We found unidirectional recruitment relations between the methylated DNA binding protein MeCP2 and KSHV encoded LANA. Where MeCP2 recruits LANA, but LANA is unable to recruit MeCP2. We were able to break this unidirectional recruitment by forcing MeCP2 dimerization. We propose that this unidirectional recruitment is the result of available interacting domains. Furthermore, this unidirectional recruitment seems to be critical for viral latency since LANA fails to maintain the viral genomes in MeCP2 mutant cells. Therefore, in this case unidirectional recruitment is a matter of survival or extinction.

## Introduction

Kaposi’s sarcoma associated herpesvirus (KSHV, HHV-8) is the causative agent of all forms of Kaposi’s sarcoma (KS) and is tightly associated with primary effusion lymphoma (PEL), and multicentric Castleman’s disease ^1,2^. Like all herpes viruses KSHV has two phases: latent (dormant) and lytic (productive) cycle. During latency the viral genomes reside in the nucleus of infected cells as large non-integrated plasmids, known as viral episomes. All KSHV infected cells express LANA, and LANA is essential for viral latency ^3–5^. LANA binding to the viral episomes is critical both for replication by recruitment of the cellular replication machinery to the viral genomes ^6,7^, and for maintenance by tethering the viral episomes to the cell chromosomes during cell division ^8–10^. Both the N and C-terminal regions of LANA have been shown to mediate chromosome association; the N-terminal via binding to core histones H2A and H2B ^11^, while for the C-terminal several candidates were identified including MeCP2, DEK and RING3 ^12–14^. Multiple LANA binding sites (LBS1, LBS2 and LBS3) ^15,16^ within the KSHV terminal repeats, and its oligomerization ability ^17,18^ create clearly visible LANA dots in KSHV-infected cells. In contrast, LANA is equally distributed in the nucleus when it is expressed in un-infected cells ^10,12,13,19^. So, it is challenging to determine whether LANA or the viral genomes are the recruiters of cellular factors.

For many nuclear processes directional recruitment of protein complexes is critical for proper function. Transcription factor that binds a specific DNA sequence recruits co-activators and general transcription factors that results in subsequent recruitment of RNA polymerase leading to gene expression. In this scenario, it is expected that a transcription factor will recruit the co-activator, and not vice versa. One example of recruitment directionality is the estrogen receptor, where the combination of ligand binding, dimerization and sequence specific DNA binding are required to generate high affinity surface to recruit co-activator proteins ^20^. This conformational change upon DNA binding and dimerization will favor binding and recruitment of co-activator to enhancer/promoter bound receptor and minimize interaction with DNA-free receptor.

MeCP2 binds methylated DNA and recruits co-repressor complexes to execute the readout of CpG methylation into transcription repression ^21^. Several studies detected interaction between MeCP2 and histone deacetylase 1 (HDAC1), and recruitment of histone deacetylase activity plays a role in transcription repression imposed by MeCP2 ^22,23^. MeCP2 and HDAC1 represent an example of two proteins that interact in the nucleus. While MeCP2 recruits HDAC1 to methylated regions in order to repress transcription, we expect that HDAC1 should not recruit MeCP2 to promoters in those cases where HDAC1 is recruited by other transcription factors. Another protein that plays important role in transcription repression is heterochromatin protein 1 (HP1). HP1 interacts with the repressive mark histone 3 lysine 9 tri-methylation (H3K9Me3) and via recruitment of SUV39H1, the histone H3K9Me3 methyltransferase, leads to spreading of this heterochromatin repressive histone mark ^24,25^. HP1 has been shown to interact with MeCP2 ^26,27^, but whether HP1 can recruit MeCP2 is an open question.

Cas9 is an endonuclease that can be directed to different DNA targets that are complimentary to a guide RNA and contain a PAM sequence ^28^. Soon after the discovery of this property, many applications of this sequence specific targeting of Cas9 emerged. For targeting purposes, the endonuclease is no longer needed, and a catalytically dead mutant Cas9 (dCas9) that still retains DNA binding capability was created ^28,29^. Targeting this dCas9 to promoter sequences results in transcription activation or repression depending on the fused domains and the targeting location relative to the transcription start site ^29,30^. Shortly after the creation of the dCas9 it was directed to repeat elements for the visualization of these elements in fixed ^31^ and live cells ^32–34^.

Here we harness CRISPR/Cas9 to determine recruitment relations between LANA and cellular proteins. We termed this method CRISPR-PITA, for Protein Interaction and Telomere recruitment Assay. We combined the ability of the SunTag system ^35^ to gather up to ten molecules of a protein of interest with CRISPR/dCas9 targeted to a repeat sequence such as telomeres. The ability of this protein to recruit endogenous or fluorescently tagged proteins to these telomere dots is determined. Interestingly, we find that the recruitment does not always work bi-directionally. LANA fails to recruit MeCP2, but MeCP2 efficiently recruits LANA. Similarly, HDAC1 that interacts with the transcriptional-repression domain (TRD) of MeCP2, same as LANA, was unable to recruit MeCP2. While HP1α that interacts with the N-terminal of MeCP2, was able to recruit MeCP2. We hypothesized that the interaction domain with LANA will become available for interaction only upon DNA binding and dimerization. Indeed, we found that forcing MeCP2 dimerization via the tandem epitopes of SunTag, enables LANA to recruit MeCP2 onto the viral genomes. The unidirectional recruitment of LANA by MeCP2 raised the possibility that MeCP2 may serve as a protein anchor for LANA tethering. Indeed, we found that the virus fails to maintain its episomes in cells expressing a mutant MeCP2 or in cells in which MeCP2 was knock-down. Therefore, the unidirectional recruitment of LANA by MeCP2 is essential for viral genome maintenance during latency.

## Results

### Creation of visible dots at the telomeres with dCas9-SunTag and scFv-LANA

The KSHV encoded LANA localizes in large dots together with the viral episomes in KSHV infected cells but is equally distributed in the nucleus when expressed in un-infected cells ^10,12,13,19^. In order to test recruitment directionality between LANA and nuclear proteins we were interested to create LANA dots in un-infected cells. The ability of dCas9 to direct proteins of interest to any genomic locus makes it an ideal system for a recruitment assay. To enhance dot formation, we utilized the SunTag system where tandem epitopes can concentrate multiple molecules of an antibody-fused protein that recognizes this epitope (up-to 10 with the tag used in this study) ^35^. In our case the tandem epitopes are fused to dCas9 and the antibody scFv is fused to our protein of interest (**Fig. 1a**). Transfection of LANA fused with the antibody scFv (scFv-LANA) together with dCas9 and sgRNA for telomere repeat sequence, resulted in LANA-telomere-dots that were visible by immuno-fluorescent assay (IFA) for LANA (**Fig. 1b**). When the dCas9 was omitted from the mix LANA could not generate nuclear dots, suggesting that LANA-telomere-dots are dependent on the targeting of LANA to telomeres via dCas9. To validate that LANA was targeted to the telomeres we used an antibody against the telomeric repeat factor 2 (TRF2), and indeed scFv-LANA co-localized with TRF2 when co-transfected with dCas9 and sgTelomere (**Fig. S1**). The ability of LANA to oligomerize ^15,17,18,36,37^, may explain the large dots we observed when we targeted LANA to telomeres. Studies of the LANA oligomerization property identified a LANA oligomerization mutant ^17,18^. Therefore, we created this LANA oligomerization mutant (F1037A/F1041A) in the context of scFv-LANA (scFv-LANA-mutant) and compared the dot size to scFv-LANA. As expected, the LANA oligomerization mutant created larger number of small telomere dots compared to wt LANA for both LANA and TRF2 (**Fig. S1**). These experiments indicate that we can generate LANA dots in un-infected cells using the dCas9-SunTag system.

**Figure 1.**
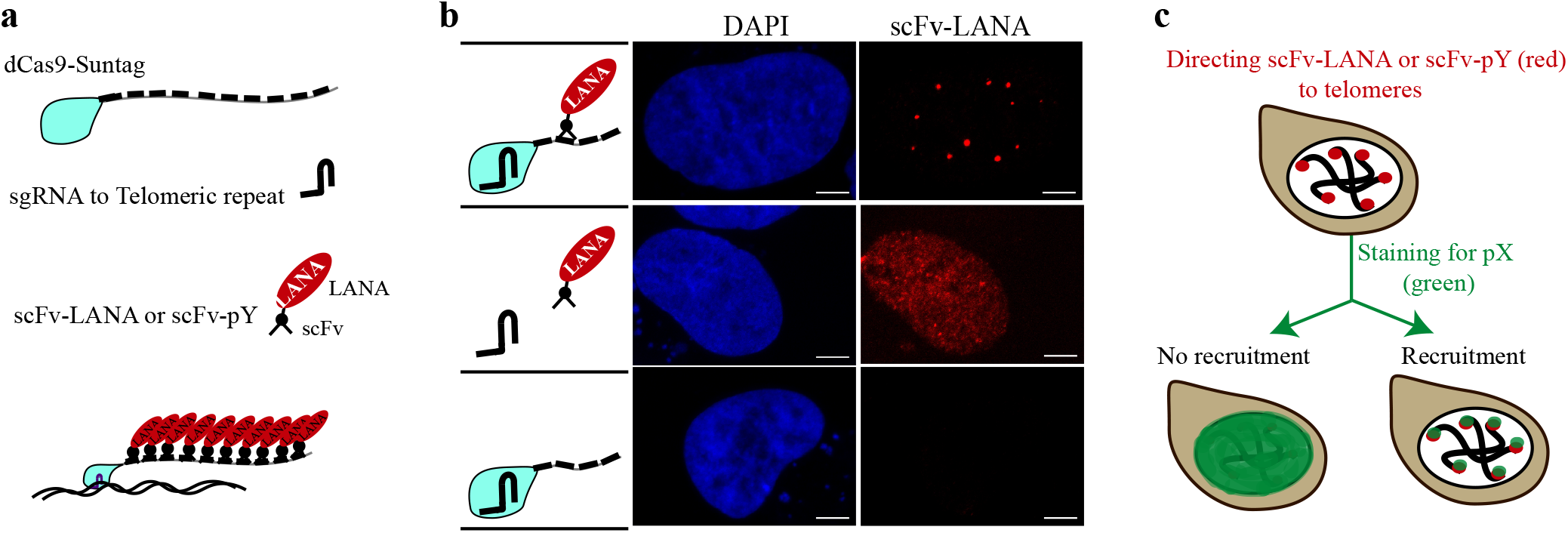
Schematic illustration of CRISPR-PITA. **(A)** Schematic illustration of dCas9-SunTag, sgRNA and scFv-LANA or other protein of interest (scFv-pY). **(B)** SLK cells were transfected with dCas9-SunTag, scFv-LANA, and sgTelomere as illustrated on the left of the images. Immunofluorescence assay was performed to detect LANA (red), and the nucleus was stained with DAPI (blue). Scale bar = 5 μm. Images are representatives of at least three independent experiments. **(C)** Schematic illustration of the CRISPR-PITA, targeting of scFv-LANA or any other protein of interest to the telomeres via dCas9. Then the recruitment of other nuclear proteins to these dots is evaluated via immunostaining.

### LANA recruits ORC2 and SIN3A but not MeCP2

In this assay a protein of interest is targeted to telomeres via dCas9-SunTag, and the recruitment of endogenous proteins to these dots is detected by immunofluorescence assay (**Fig. 1c**). We termed this method CRISPR-PITA for Protein Interaction and Telomere recruitment Assay. We performed the CRISPR-PITA assay for ORC2, a subunit of the Origin Replication Complex (ORC), previously reported to interact with LANA and localize with LANA-episome-dots ^18,38–40^. ORC2 was recruited by LANA, as was determined by the easily detected dots that co-localized with LANA (**Fig. 2a**), and the overlapping fluorescence intensity peaks along the arrow line (on the right side of the images). It is important to note, that while the dCas9-SunTag and scFv-LANA were ectopically expressed, the recruited proteins are endogenous proteins, indicating the ability of LANA to recruit the endogenous ORC2.

**Figure 2.**
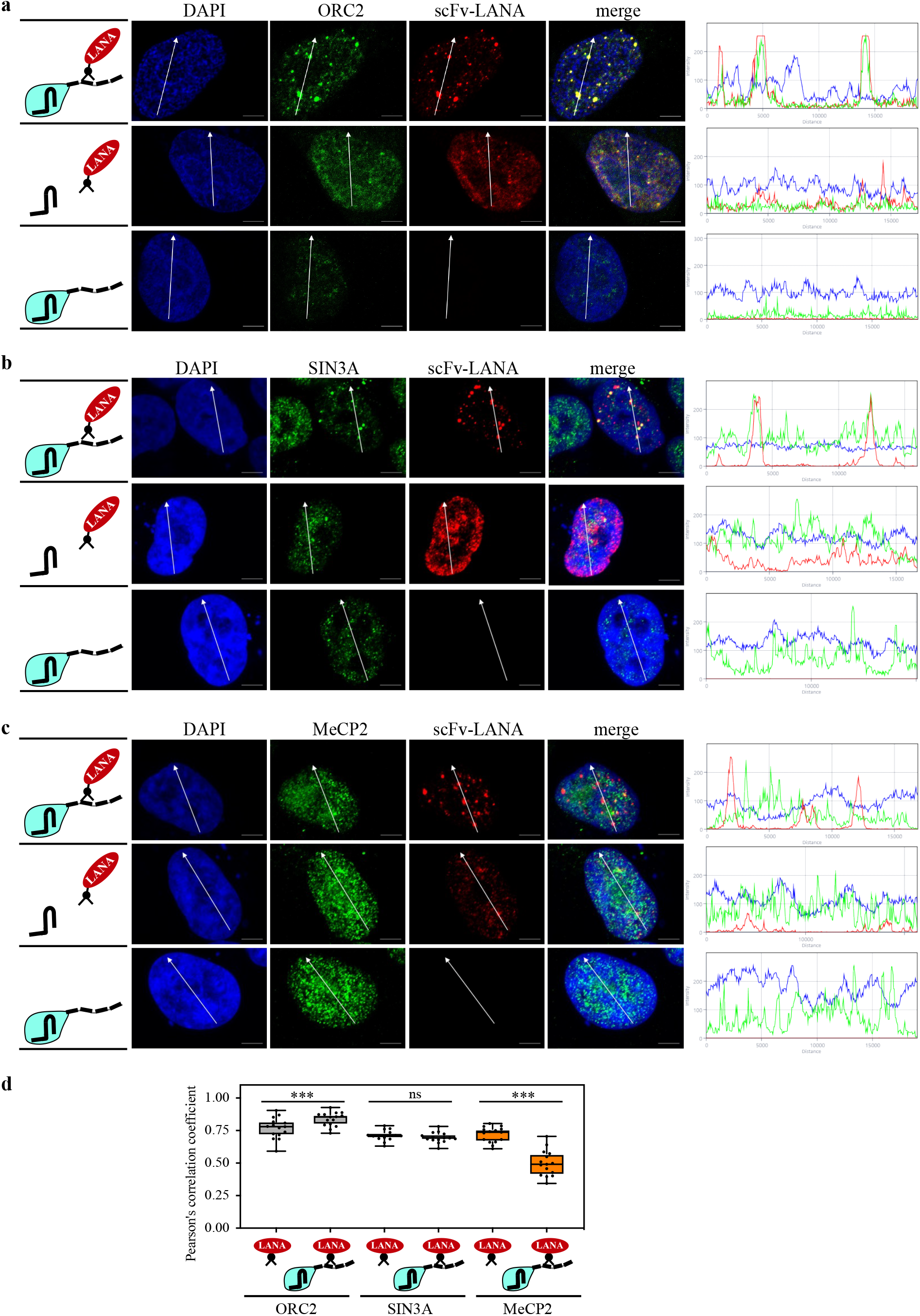
LANA recruits ORC2 and SIN3A but not MeCP2. SLK cells were transfected with dCas9-SunTag, scFv-LANA, and sgTelomere expression plasmids as can be seen in the illustrations on the left. Immunofluorescence assays were performed to detect LANA (red) and ORC2 **(A)** mSin3a **(B)** or MeCP2 **(C)** (in green) cellular localization. The nucleus was stained with DAPI. Scale bar = 5 μm. Images are representatives of at least three independent experiments. The plots of the red, green and blue pixel intensities along the white arrow (in the left panels) are presented. (**D**) Pearson’s correlation coefficient was determined by ImageJ (JACoP Pluing) ^74^ for 15 different cells in each treatment, and presented as box and whiskers (min to max). Two-tailed *t* tests were performed (*, P ≤ 0.05; **, P ≤ 0.01; ***; P ≤ 0.001).

Both the corepressor protein SIN3A (mSin3a) ^40,41^, and the methyl-CpG binding protein MeCP2 ^12,13,42^, have been reported to interact with LANA. However, MeCP2 seems not to be associated with LANA dots in KSHV infected cells ^43^. We found that scFv-LANA recruited SIN3A to LANA-telomere-dots, both by the visual observation of the dots and the fluorescence intensity plot (**Fig. 2b**). Analysis of MeCP2 recruitment via the CRISPR-PITA revealed no recruitment of MeCP2 to LANA-telomere-dots (**Fig. 2c**). We also determined the Pearson’s correlation coefficient for the different treatments (**Fig. 2d**). Since these proteins are known LANA interactors, we expected to find overlap when LANA alone (without dCas9) is transfected. When LANA is targeted to the telomeres via dCas9, if LANA is able to recruit its partner, then the overlap is expected to remain or even increase. In contrast, if LANA is unable to recruit its partner, the overlap should be reduced. This is exactly what we found when we measured the Pearson’s correlation coefficient for 15 cells in each experiment. The overlap was reduced for the non-recruited MeCP2, remained the same for SIN3A, and increased for ORC2.

One possibility to explain the lack of MeCP2 recruitment by LANA, is that as a methylated DNA binding protein, it is engaged in binding methylated DNA and therefore not available to be recruited by LANA. To test this, we performed the CRISPR-PITA in cells with knock-out (KO) of DNMT1 and DNMT3b (HCT DKO) that were reported to lose over 95% of their CpG methylation ^44^. Even in HCT DKO, scFv-LANA could not recruit MeCP2 to LANA-telomere-dots (**Fig. S2**), suggesting that protein availability could not explain the lack of recruitment in this case. Immunofluorescence assays for ORC2, SIN3A and MeCP2 in KSHV-infected PEL (BCBL1) cells, revealed that while ORC2 and SIN3A were found associated with LANA-episome-dots, MeCP2 was not associated with the LANA-episome-dots (**Fig. S3**), supporting the ability of the CRISPR-PITA to predict recruitment ability between proteins.

### MeCP2 recruits LANA

The observation that scFv-LANA could not recruit MeCP2, despite their documented interaction ^12,13,42^ highlights the importance to determine recruitment relations between proteins. Krithivas et al. have shown that expression of human MeCP2 is able to recruit LANA to heterochromatin in NIH 3T3 mouse cells ^13^. Therefore, we tested the ability of scFv-MeCP2 to recruit LANA to MeCP2-telomere-dots. Cells were co-transfected with dCas9, sgTelomere, scFv-MeCP2 and expression vectors for GFP-LANA or GFP alone as control. The scFv-MeCP2 was able to recruit GFP-LANA to MeCP2-telomere-dots, but not GFP alone, supporting the ability of MeCP2 to recruit LANA (**Fig. 3a,b**). LANA contains three distinct domains, the N-terminal region (AA 1-329), the C-terminal (AA 936-1162), and a middle repeat region. Both the N and C-terminal regions have been shown to mediate chromosome association; the N-terminal via binding to core histones H2A and H2B ^11^, while for the C-terminal several candidates were identified including MeCP2, DEK and RING3 ^12–14^. To determine the region of LANA sufficient for recruitment by MeCP2, we performed CRISPR-PITA by scFv-MeCP2 for LANA N+C, LANA N and LANA C. While both LANA N+C and LANA C were efficiently recruited by MeCP2, no recruitment was observed with the N-terminal (**Fig. 3c,d**). This result agrees with a previous study that located the interaction between LANA and MeCP2 to the C-terminal region ^12^. Importantly, this result further supports the notion that despite the high concentration of the recruiter protein at telomeres, still CRISPR-PITA requires specific interaction, as both the N-terminal of LANA and GFP-alone were not recruited by MeCP2 in our assay.

**Figure 3.**
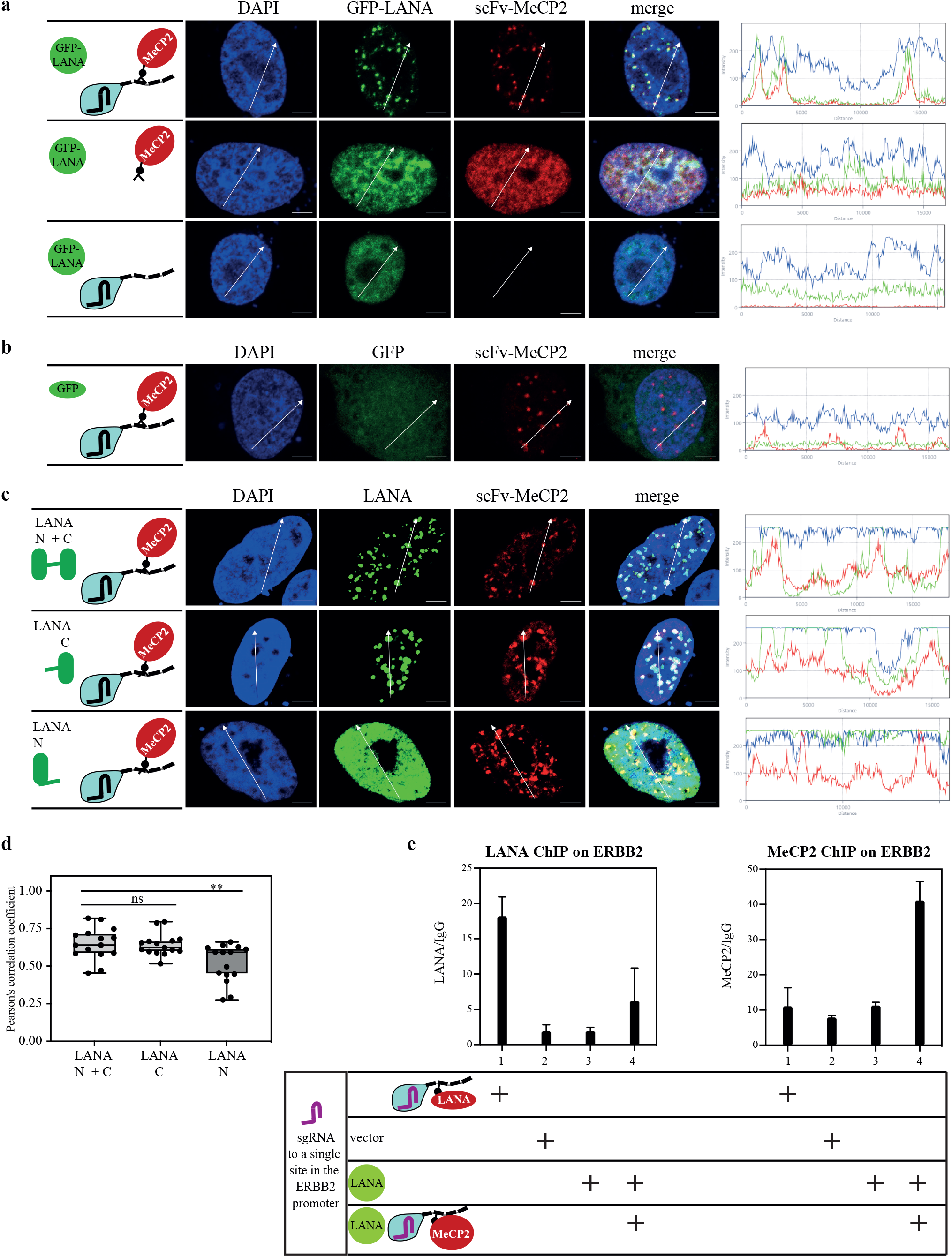
MeCP2 recruits LANA. SLK cells were transfected with dCas9-SunTag, scFv-MeCP2, and sgTelomere in combination with GFP-LANA **(A)** or GFP **(B)** as illustrated on the left. Immunofluorescence assay was performed to detect scFv-MeCP2, or fluorescently labelled GFP-LANA and GFP. The nucleus was stained with DAPI. Scale bar = 5 μm. Images are representatives of at least three independent experiments. The plots of the red, green and blue pixel intensities along the white arrow (in the left panels) are presented. (**C**) SLK cells were transfected with FLAG-LANA N+C, FLAG-LANA C, or FLAG-LANA N, same as in A. Immunofluorescence assay was performed to detect both scFv-MeCP2 and LANA. (**D**) Pearson’s correlation coefficient was determined for 15 different cells in each treatment, and presented as box and whiskers (min to max). Two-tailed *t* tests were performed (*, P ≤ 0.05; **, P ≤ 0.01; ***; P ≤ 0.001). (**E**) ChIP assay to detect LANA (left panel) and MeCP2 (right panel) association with the ERBB2 promoter following recruitment with sgRNA to this locus.

Targeting dCas9 to a repeat element, results in high protein concentration at specific nuclear locations. One concern that might be raised is whether the recruitment we observed with CRISPR-PITA is specific or an artifact of the high protein concentration. To rule out this possibility, we directed dCas9 to a single chromosomal location (at ERBB2 promoter) and followed the association of the proteins with this location by chromatin immunoprecipitation (ChIP) assay followed by qPCR (**Fig. 3e**). Here again we found that scFv-MeCP2 recruited more LANA to this promoter, while scFv-LANA failed to do the same for MeCP2. This result indicates that despite targeting dCas9 to a repeat sequence, the recruitment in CRISPR-PITA nicely reflects the recruitment by a single locus. This set of experiments highlights the distinction between interaction and recruitment, while LANA and MeCP2 were shown to interact with each other, still MeCP2 can recruit LANA, but LANA cannot recruit MeCP2.

### MeCP2 recruits HDAC1, but HDAC1 is unable to recruit MeCP2

Several studies detected interaction between MeCP2 and histone deacetylase 1 (HDAC1), and this recruitment of histone deacetylase activity plays a role in transcription repression imposed by MeCP2 ^22,23^. MeCP2 and HDAC1 represent an example where two proteins can interact to form protein complexes in the nucleus. We expect that MeCP2 should recruit HDAC1 to methylated regions to repress transcription, but HDAC1 should not recruit MeCP2 to promoters where it is recruited by other transcription factors. Therefore, we applied the CRISPR-PITA to determine recruitment relations between these two cellular proteins. We found that scFv-MeCP2 efficiently recruits HDAC1 (**Fig. 4a**). In contrast, scFv-HDAC1 failed to recruit MeCP2 (**Fig. 4b,c**). Thus, in the context of MeCP2 and HDAC1, recruitment is also unidirectional (**Fig. 4d**), enabling MeCP2 to recruit HDAC1 to repress transcription, but preventing the mislocalization of MeCP2 by HDAC1.

**Figure 4.**
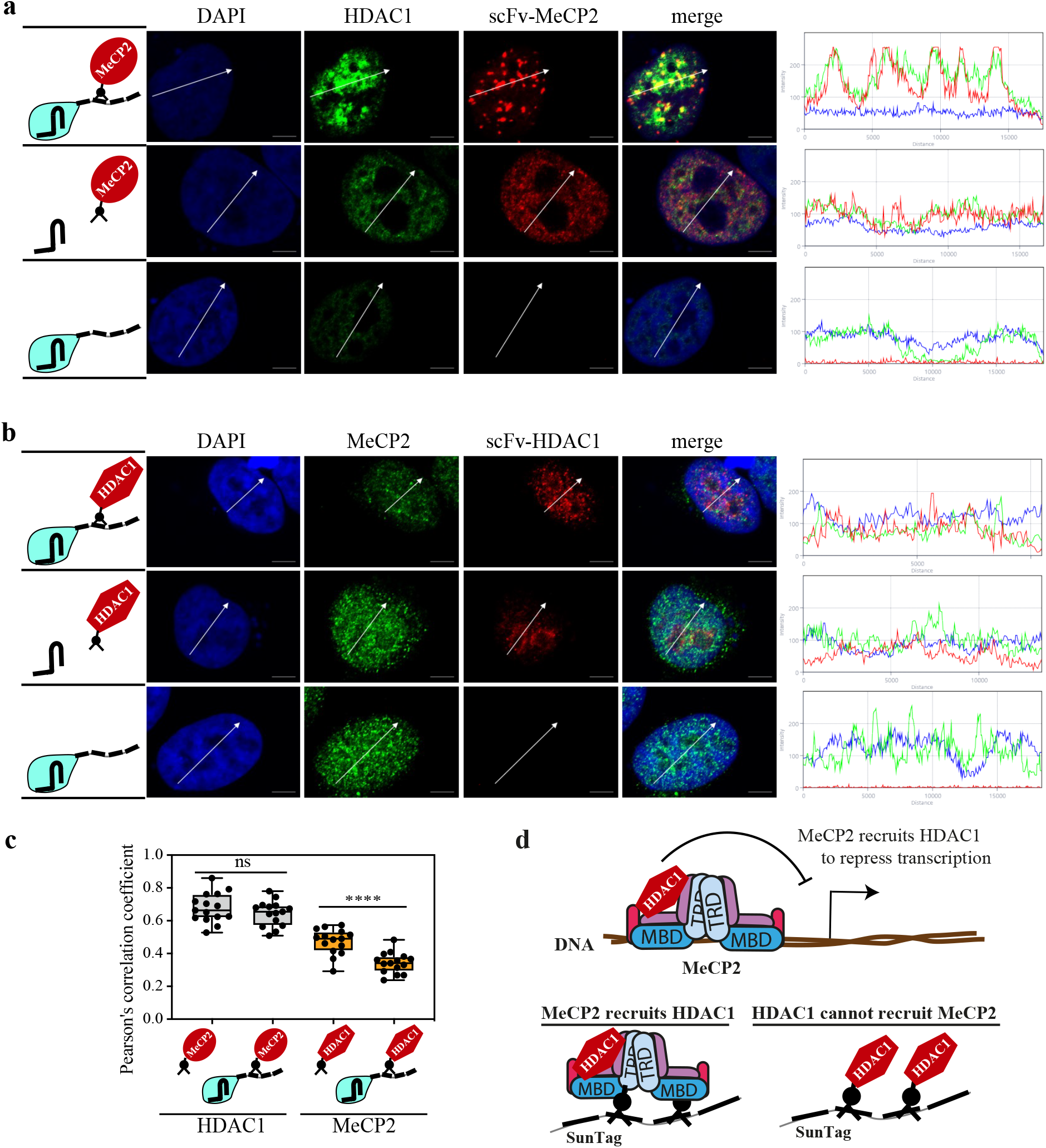
MeCP2 recruits HDAC1, while HDAC1 cannot recruit MeCP2. **(A)** SLK cells were transfected with dCas9-SunTag, scFv-MeCP2 and sgTelomere as illustrated on the left. Immunofluorescence assay was performed to detect HDAC1 (green) and scFv-MeCP2 (red). **(B)** SLK cells were transfected same as in A but with scFv-HDAC1. Immunofluorescence assay was performed to detect MeCP2 (green) and scFv-HDAC1 (red). The nucleus was stained with DAPI. Scale bar = 5 μm. Images are representatives of at least two independent experiments. The plots of the red, green and blue pixel intensities along the white arrow (in the left panels) are presented. **(C)** Pearson’s correlation coefficient was determined for 15 cells in each treatment and presented as box and whiskers (min to max). Two-tailed *t* tests were performed (****; P ≤ 0.0001). (**D**) Schematic illustrations of the current model where MeCP2 recruits HDAC1 to repress transcription, and this model is supported in CRISPR-PITA that MeCP2 recruits HDAC1 but HDAC1 cannot recruit MeCP2.

### Availability of interacting domains determines recruitment directionality

A unidirectional recruitment between two interacting proteins is achieved when the interacting domain is available only upon certain conditions. In the case of MeCP2, it is expected that the transcriptional-repression domain (TRD) will be unavailable when it is a free protein in solution and becomes available for protein interaction only when MeCP2 dimerizes upon binding to DNA (**Fig. 5d,e**). Both LANA and HDAC1 interact with the same region of MeCP2, the transcriptional-repression domain (TRD) and the methyl-CpG-binding domain (MBD) ^36,41^. In contrast, Heterochromatin protein 1 (HP1α) interacts with the N-terminal domain (amino acids 1–55) of MeCP2 ^26^ (**Fig. 5a**). We hypothesized that since HP1α interacts with a different domain that might be available for interaction also in its DNA-free state, HP1α might be able to recruit MeCP2. Indeed, we found that scFv-HP1α was able to recruit MeCP2 in CRISPR-PITA (**Fig. 5b,c**). This result supports the notion that available interacting domains determine recruitment directionality (**Fig. 5d,e).** A previous study ^43^, and our result (**Fig. S3**), show that LANA fails to recruit MeCP2 to LANA episome dots in infected cells. We hypothesized that forcing dimerization to DNA-free MeCP2 will break the unidirectional recruitment and enable LANA to recruit MeCP2. Although CRISPR-PITA does not directly test the DNA binding of MeCP2, however the tandem epitopes SunTag brings several MeCP2 molecules together and therefore might force dimerization and by that can mimic DNA bound MeCP2 (**Fig. 5f**). KSHV infected cells were transfected with scFv-MeCP2 with or without SunTag, to force MeCP2 dimerization. To exclude the possibility that the dCas9 attached to the SunTag will bind DNA even without sgRNA, we expressed the tandem epitopes SunTag only, without dCas9. Co-transfection of scFv-MeCP2 together with a SunTag resulted in recruitment of scFv-MeCP2 to LANA dots in infected cells (**Fig. 5g-i**). While transfection of only scFv-MeCP2 could not recruit MeCP2 to LANA dots as expected. Our result that we can break the unidirectional recruitment relations between LANA and MeCP2, by forcing MeCP2 dimerization supports our model of available interacting domains.

**Figure 5.**
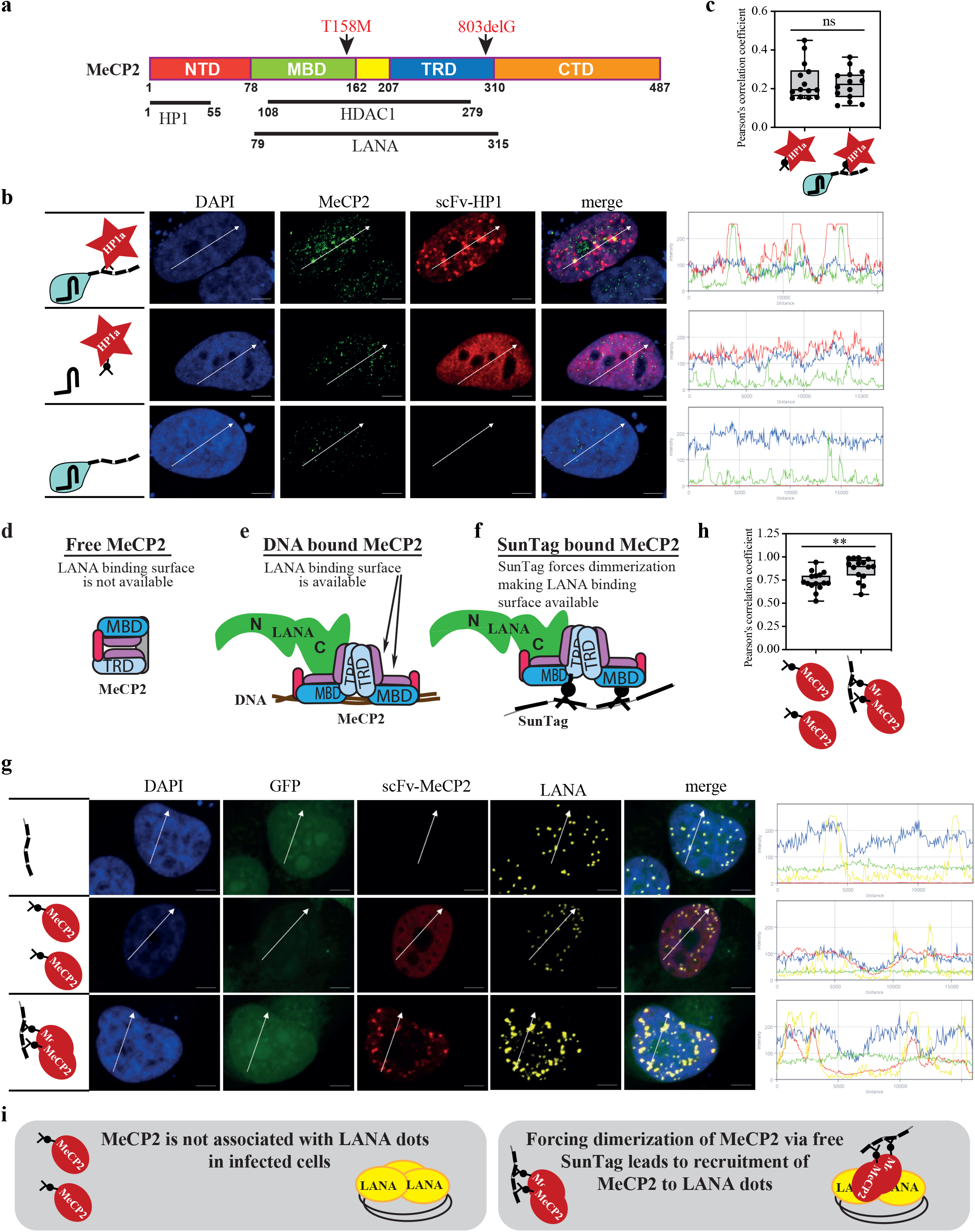
Availability of interacting domains determines recruitment directionality. (**A**) schematic presentation of MeCP2 and the interaction domains with LANA, HDAC1, and HP1α. Two Rett syndrome mutants are indicated above. (**B**) SLK cells were transfected with dCas9-SunTag, scFv-HP1α, and sgTelomere as illustrated on the left. Immunofluorescence assay was performed to detect MeCP2 (green) and scFv-HP1α (red). The nucleus was stained with DAPI. Scale bar = 5 μm. Images are representatives of at least two independent experiments. (**C**) Pearson’s correlation coefficient was determined for 15 cells in each treatment and presented as box and whiskers (min to max). Two-tailed *t* tests were performed. Schematic illustrations of the model for free (**D**) DNA bound MeCP2 (**E**) and SunTag scFv-MeCP2 (**F**) are presented. (**G**) iSLK infected cells (iSLK.Bac16) were transfected with free SunTag and scFv-MeCP2. Then, immunofluorescence assay was performed to detect scFv-MeCP2 (red) and LANA (yellow). GFP is a marker of infected cells. The plots of the red, yellow, green and blue pixel intensities along the white arrow (in the left panels) are presented. (**H**) Pearson’s correlation coefficient was determined for 15 cells in each treatment, and presented as box and whiskers (min to max). (**H**) Schematic presentation of the experiment in G, and its conclusions.

### knock-down of MeCP2 reduces KSHV genome maintenance during latency

One important function of LANA is to maintain the viral episomal genomes, by tethering the viral genomes to cellular chromosomes during cell division, ensuring they remain in the nucleus once nuclear envelope is reformed. To do so, LANA directly binds the LANA binding sites (LBS1, LBS2 and LBS3) ^15,16^ on the viral genomes, and associates with cellular chromosomes. Our finding of the unidirectional recruitment relations between MeCP2 and LANA, suggests that the interaction with MeCP2 is permitted only once it is bound to methylated DNA, and therefore may be a potential candidate for LANA tethering. To test this possibility, we knock-down (KD) MeCP2 and followed the ability of KSHV to maintain latency. SLK cells infected with a recombinant KSHV virus Bac16 (SLK.Bac16) ^45^ that confers hygromycin resistance were transduced with lentivirus expressing MeCP2 shRNA or control shRNA. Cells were grown in the presence of hygromycin and the number of colonies were counted (**Fig. 6a**). The KD of MeCP2 was verified by RT-qPCR (**Fig. 6b**). We found that KD of MeCP2 dramatically reduced the ability of KSHV to maintain latency and to enable the growth of cells in the presence of hygromycin (**Fig. 6c,d**).

**Figure 6.**
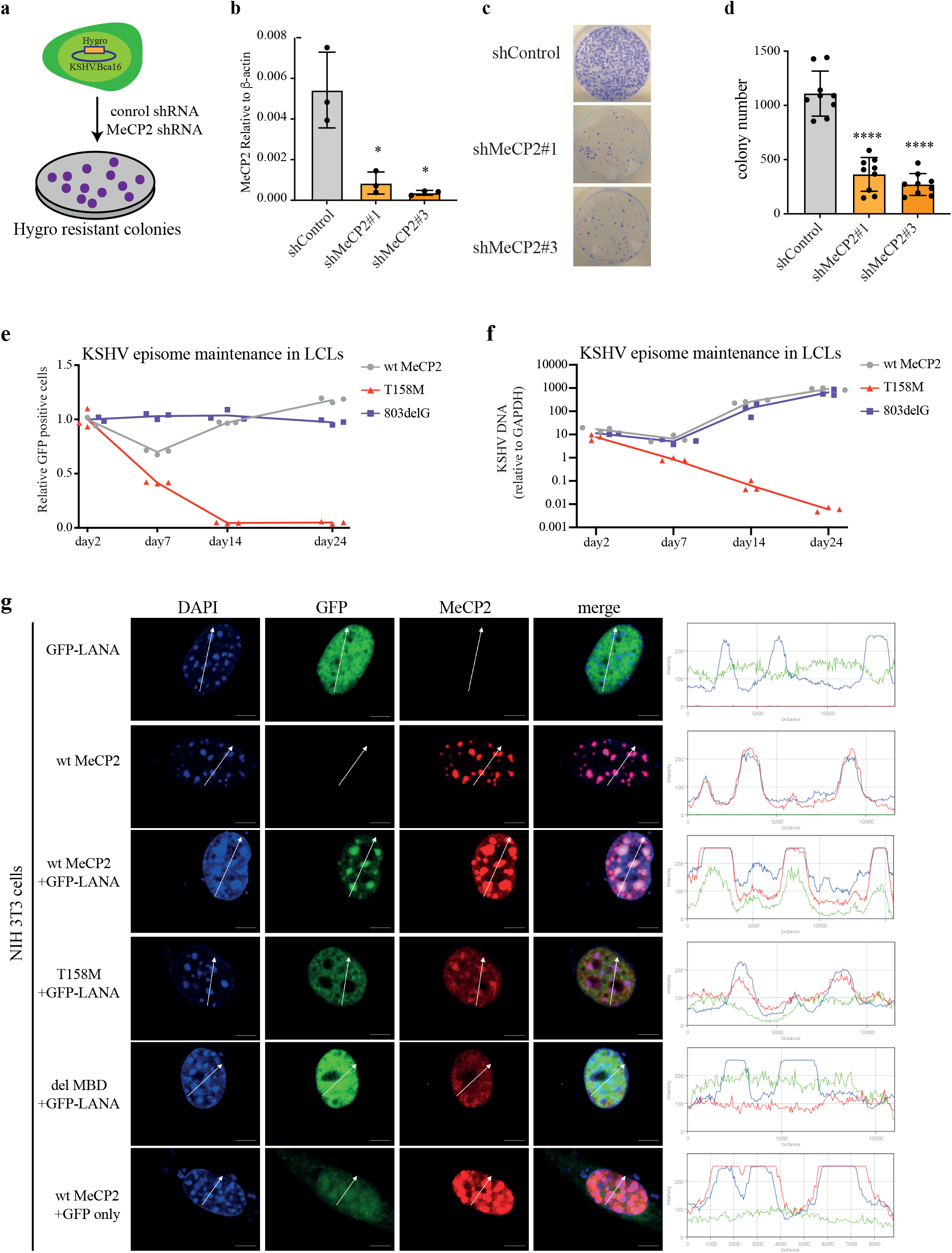
MeCP2 is essential for viral episome maintenance. **(A-D)** shRNA against MeCP2 reduces KSHV latency. SLK cells infected with rKSHV-Bac16 (SLK.Bac16) were transduced with lentivirus expressing shRNA against MeCP2 or control (**A**) and the expression level of MeCP2 was determined by RT-qPCR (**B**). The cells were plated in the presence of hygromycin to select for latently infected cells. Representative pictures of such plates (**C**) and summary of colony number in three independent experiments, each with three biological repeats (**D**). **(E-F)** Rett syndrome mutant MeCP2 cannot support KSHV maintenance during latency. LCLs derived from Rett syndrome patients expressing a wt or mutant MeCP2, were infected with rKSHV.219 and GFP-positive cells were quantified by FACS analysis (**E**) or amount of KSHV DNA by qPCR (**F**) were evaluated at the indicated time points. Experiments were performed with biological triplicates. (**G**) NIH 3T3 cells were transfected with GFP-LANA and wt MeCP2, T158M mutant, or del-MBD. Immunofluorescence assay was performed to detect LANA (green) and MeCP2 (red). The heterochromatin foci in the nucleus of NIH3T3 mouse cells has high intensity of DAPI (blue). Scale bar = 5 μm. Images are representatives of at least two independent experiments. The plots of the red, green and blue pixel intensities along the white arrow (in the left panels) are presented.

### Rett syndrome mutant MeCP2 cannot support KSHV maintenance during latency

Sporadic mutations in MeCP2 are associated with most cases of Rett syndrome (RTT), a severe X-linked neurodevelopmental disorder in humans ^46^. Lymphoblastoid Cell Lines (LCLs) derived from RTT patients ^47^ expressing wild-type (wt), mutant T158M (that cannot bind methylated DNA), or 803delG (deletion of the C’ terminus) of MeCP2 (**Fig. 5a**), were tested for their ability to support KSHV episome maintenance. Cells were infected with rKSHV (rKSHV.219) expressing GFP and maintenance of the viral episomal DNA and the percentages of GFP positive cells were analyzed (**Fig. 6e,f**). While both wt and 803delG efficiently supported viral latency, the MeCP2 mutant in the MBD that cannot bind DNA (T158M) could not. This result indicates that a functional MeCP2 that is able to bind methylated DNA is essential for KSHV genome maintenance. The fact that KSHV can maintain latency without the need for drug selection in LCLs, can be explained by the co-infection of KSHV and Epstein-Barr virus (EBV) in these cells. A previous study already shown that EBV can enhance KSHV maintenance ^48^. LANA has been shown to bind both the MBD and TRD of MeCP2 ^12^. To determine if LANA can be recruited by the T158M mutant, we cloned it into scFv-MeCP2 and performed CRISPR-PITA assay. We found that MeCP2 T158M mutant can recruit LANA, when it is artificially targeted to chromatin by dCas9 and forced to dimerize via the SunTag (**Fig. S4**). While the MBD deletion mutant, lost the ability to recruit LANA when artificially targeted to chromatin. To assay for both abilities, to recruit LANA and to bind methylated DNA, we harnessed a specific feature of MeCP2 to localize to heterochromatin loci in mouse cells. This assay is typically performed in NIH 3T3 cells, since they express very low levels of endogenous MeCP2 ^13,49^. Similar to previous studies ^12,13^, we found that LANA can be efficiently recruited to heterochromatin when co-transfected with wt MeCP2 (**Fig. 6g**). In contrast, both T158M and MBD-del mutants could not recruit LANA to heterochromatin loci. Altogether, we found that Rett syndrome mutant MeCP2 T158M, that cannot recruit LANA to methylated DNA, also cannot support KSHV episome maintenance. We revealed a critical role for MeCP2 in KSHV episome maintenance by LANA.

## Discussion

A powerful method to study recruitment is the Chromatin immunoprecipitation (ChIP) assay, that can detect association of proteins with specific DNA sequences ^50,51^. ChIP assays require fixation, proper shearing of the chromatin, and special antibodies that can perform the immunoprecipitation of the cross-linked chromatin. In many cases even when two or more proteins are associated with the same genomic region, it is hard to evaluate the relationship between the proteins and determine essential factors for the recruitment. Although this limitation can be overcome by using Knockout (KO) or Knockdown (KD) of one protein and testing the ability of other proteins to be recruited to DNA/chromatin in the absence of that protein, this raises additional challenges in creating KO or KD of the target gene. Another recruitment assay is based on integration of the *lac*O array into the genome and directing the protein of interest to this array via fusion to the Lac-repressor ^52^. One limitation of this assay is the limited number of cell lines that were generated to contain integration of this *lac*O array. In a previous study, KSHV encoded LANA was directed to the host genome via the *lac*O array ^53^. In this study LANA was able to recruit the two bromo and extra terminal domain (BET) proteins, BRD2 and BRD4, two known LANA interacting proteins. However, because of the system limitation, the ability of BRD2 and BRD4 to recruit LANA was not tested.

Here we describe a simple method to test for recruitment relations between proteins. This method does not require chromatin shearing and can be performed with various antibodies that can recognize the native form of proteins, and in the absence of such an antibody a recruitment of fluorescently labeled proteins can be observed. To obtain visible dots we combined the ability of the repeated epitope SunTag with dCas9 targeted to a repetitive genomic sequence such as the telomeres. Although targeting dCas9 to telomeres has been performed previously to follow nuclear events such as, liquid–liquid phase separation (LLPS), or the generation of heterochromatin ^54,55^, but has not been applied to test recruitment directionality. Using this method, we found unidirectional recruitment relations between proteins. Our results with CRISPR-PITA, indicated that LANA was able to recruit ORC2 and SIN3A, but was unable to recruit MeCP2 to LANA-telomere-dots. While it has been shown that ORC2 localizes with LANA dots ^18,38–40^, and MeCP2 does not ^43^, there was no published data on SIN3A localization in KSHV infected cells. So, we performed immunofluorescence assay in KSHV-infected BCBL1 cells and found that both ORC2 and SIN3A were associated with LANA dots, but MeCP2 was not. The immunofluorescence assays in KSHV-infected cells agree with the CRISPR-PITA results, supporting the ability of CRISPR-PITA to predict recruitment ability of proteins. While both ORC2 and SIN3A were recruited by LANA in CRISPR-PITA, it seems that ORC2 was recruited to most LANA dots, while SIN3A was associated only with part of LANA dots. This difference was also reflected in the Pearson’s correlation coefficient, where the correlation with SIN3A remained the same while the correlation with ORC2 increased upon directing LANA to telomeres. Interestingly, this difference was also observed in infected cells, where SIN3A seems to colocalize with only part of LANA-episome dots. The recruitment of GFP-LANA by scFv-MeCP2 also suggests that CRISPR-PITA can be done without cell fixation or immunofluorescence, but directly via detection of fluorescently-labeled proteins. This property might be more relevant in cases where CRISPR-PITA is applied for screening purposes.

Targeting dCas9 to a repeat element, results in high protein concentration at specific nuclear locations. One concern that might be raised is whether the recruitment we observed with CRISPR-PITA is specific or an artifact of the high protein concentration. Our result that the same recruitment relations are detected also when we directed dCas9 to a single locus (ERBB2 promoter), rules out this possibility. Regarding specificity, we found that scFv-MeCP2 recruited the C-terminal of LANA, but not its N-terminal. Interaction of LANA C-terminal domain with MeCP2 is in agreement with a previous study for LANA ^12^. The related gamma-2 herpesvirus, herpesvirus saimiri (HVS) encodes for ORF73, a functional homolog of KSHV encoded LANA (also known as ORF73). Interestingly, also HVS encoded ORF73 interacts with MeCP2 via its C-terminal domain ^49^. The lack of recruitment for the N-terminal domain of LANA, indicated that recruitment is still specific and requires the interaction domain. Another support for the specific recruitment by CRISPR-PITA is the lack of recruitment observed for GFP alone. A growing list of nuclear proteins with intrinsically disordered regions and RNA/DNA binding properties can promote liquid–liquid phase separation (LLPS); both LANA and MeCP2 possess this property ^56,57^. Rett syndrome MeCP2 mutants, including T158M are impaired in LLPS formation. Since T158M mutant still can recruit LANA in CRISPR-PITA, excludes the possibility that recruitment in CRISPR-PITA is dependent on LLPS formation.

The inability of LANA to recruit MeCP2, despite their documented interaction prompted us to test whether MeCP2 can recruit LANA. Indeed, in CRISPR-PITA scFv-MeCP2 was able to recruit LANA. This result comes in agreement with a previous study where MeCP2 recruited LANA to mouse heterochromatin foci ^13^. This result also highlights the importance of a recruitment assay as recruitment is not always bi-directional, and in many cases should be functionally unidirectional. One of LANA functions where unidirectional recruitment seems essential, is LANA tethering of the viral episomal genome to the cell chromosomes during cell division. Therefore, we tested the possible role of MeCP2 in KSHV episome maintenance. We found that Rett syndrome mutant MeCP2, that cannot bind methylated DNA (T158M), could not support KSHV latency. In addition, KSHV failed to maintain its genomes following MeCP2 KD. Our results suggest that in addition to LANA interaction with histones ^11^, MeCP2 adds another layer to secure chromosome tethering (suggested model in **Fig. S5**). Kelley-Clarke et al. ^8^ have found that LANA with a reduced histone binding due to N-terminus mutation, is dependent on the C-terminus for mitotic chromosome binding and episome persistence. Similar to KSHV encoded LANA, HVS encoded ORF73 has a critical role in viral episomal genome maintenance^58^. MeCP2 has been shown to be essential for tethering of the viral genomes by ORF73 to maintain HVS latency ^49^. Therefore, our observation that MeCP2 is essential for episome maintenance, seems to be conserved between KSHV and HVS. This conservation for MeCP2 tethering via the C-terminus of LANA agrees also with a previous hypothesis that the C-terminal of LANA was the original chromosome tether, and the N-terminal tether evolved in a later stage ^59^. It has been shown that LANA tethers the viral episomes onto pericentromeric regions ^60,61^, and it has been suggested that the C-terminal of LANA interacts with a cellular factor that concentrates at pericentromeric regions ^61^. Interestingly, MeCP2 is known to concentrate at pericentromeric regions ^62,63^. In our proposed model, LANA/ORF73 binds MeCP2 to anchor the viral episomes to cellular chromosomes. If the anchor (MeCP2) is free (not bound to DNA/chromosome), this may lead to viral loss. Therefore, in this case unidirectional recruitment is a matter of survival or extinction.

Do Rett syndrome patients are resistant to KSHV viral latency? *MeCP2* gene is located on the X-chromosome, and Rett syndrome is found almost exclusively in females. Females, inherent two copies of X-chromosome, one maternal and one paternal. During development one X-chromosome, is subjected to X-chromosome inactivation, allowing expression of only one copy. Since females will present a mosaic expression of the mutant and wt *MeCP2* alleles in different cells, it is reasonable to speculate that female Rett syndrome patients will not be resistant to KSHV latent infection. It is important to note that in our experiments, we used Single-cell derived clonal LCLs expressing either wt or mutant Rett syndrome MeCP2 allele ^47^. Rett syndrome in males is very rare leading to in-utero lethality in most cases due to having only a single copy of non-functional MeCP2. Another factor that contributes to the rarity of Rett syndrome in males is their maternal origin of x-chromosome. While most MeCP2 mutations are sporadic, and preferentially occur on the paternal x-chromosome ^64^ ^65^. So, only in these very rare cases of male Rett syndrome, and only in those cases that the mutation disrupts DNA binding or the TRD domain of MeCP2 we can expect a resistance to KSHV latency.

A nice example of directional recruitment is the case of MeCP2 that binds methylated DNA and recruits co-repressor complexes including HDAC1 ^22,23^. In this case we expect that MeCP2 should recruit HDAC1, leading to transcription repression. On the other hand, HDAC1 is recruited by many transcription factors to generate repressive chromatin. Therefore, we expect that it should not recruit MeCP2 to all the chromosome locations it is recruited by other transcription repressors. Indeed, we found that scFv-MeCP2 was able to recruit HDAC1 to MeCP2-telomere-dots, but scFv-HDAC1 was unable to recruit MeCP2. Both LANA and HDAC1 interact with the MBD and TRD of MeCP2 ^12,23^. In contrast, HP1α interacts with the N-terminal domain (amino acids 1–55) of MeCP2 ^26^. Indeed, we found that HP1α can recruit MeCP2 to HP1α-telomere-dots. It has been shown that phosphorylation at serine

229 (pS229) of MeCP2 strongly enhances the interaction between MeCP2 and HP1α ^27^. Our observation that HP1α can recruit MeCP2, may explain their observation that MeCP2 pS229 was enriched at the receptor tyrosine kinase gene *RET* promoter. In the future, it will be interesting to test the role of HP1α in the global distribution of MeCP2.

How come two proteins that interact with each other can generate unidirectional recruitment? MeCP2 is an intrinsically disordered protein that is monomeric in solution even at high concentrations but dimerizes upon DNA binding ^66,67^. Although CRISPR-PITA does not directly test the DNA binding of MeCP2, however the SunTag brings several MeCP2 molecules together and therefore can force dimerization. This might explain how both wt and T158M mutant MeCP2 which is unable to bind DNA, are able to recruit LANA when dimerization is forced via the SunTag in CRISPR-PITA. To test this more directly, we forced MeCP2 dimerization with a free SunTag. Under these conditions LANA was able to recruit MeCP2 to LANA-episome dots in infected cells. Revealing the mechanism behind unidirectional recruitment, enabled us to break this unidirectionality. Our study and the CRISPR-PITA may lead to a deeper understanding of complex formation and recruitment directionality in many cellular and viral interactions.

## Materials and Methods

### Cell culture

SLK cells (kindly provided by Don Ganem & Rolf Renne), SLK cells infected with KSHV.Bac16 (kindly provided by Jae U. Jung), NIH 3T3 cells (kindly provided by Oren Kobiler) and Vero-rKSHV.219 cells (kindly provided by Jeffrey Vieira) were cultured in Dulbecco’s modified Eagle’s medium (DMEM) supplemented with 10% fetal bovine serum (heat inactivated) and 100 U/ml penicillin, 100 μg/ml streptomycin, 2 mM L-glutamine, and 1 mM Sodium-Pyruvate in 5% CO_2_ at 37°C. HCT and HCT Dnmt1-/- Dnmt3b -/- (DKO) cells (kindly provided by Bert Vogelstein) were grown in McCoy’s 5A medium supplemented with the above-mentioned supplements. BJAB and BCBL1 cells (kindly provided by Richard F. Ambinder), and LCLs (kindly provided by Uta Francke), were cultured in RPMI-1640 medium supplemented with 20% FBS and the above-mentioned supplements. Single-cell derived clonal LCLs expressing the wt or mutant Rett syndrome MeCP2 allele were established in the lab of Uta Francke ^47^.

### Plasmids

pHRdSV40-dCas9-10xGCN4_v4-P2A-BFP (Addgene plasmid # 60903; http://n2t.net/addgene:60903; RRID:Addgene_60903) and pHR-scFv-GCN4-sfGFP-GB1-dWPRE (Addgene plasmid # 60907; http://n2t.net/addgene:60907; RRID:Addgene_60907) were a gift from Ron Vale ^35^. pgRNA-humanized was a gift from Stanley Qi (Addgene plasmid # 44248; http://n2t.net/addgene:44248; RRID:Addgene_44248) ^29^. The sgRNA targeting telomeres was cloned into pgRNA-humanized. The mCherry in the pgRNA-humanized was deleted by restriction digestion and ligation. LANA, HDAC1, HP1α and MeCP2 were PCR amplified and cloned into pHR-scFv-GCN4-sfGFP-GB1-dWPRE. scFv-LANA was further subcloned into pLIX_403, a gift from David Root (Addgene plasmid # 41395; http://n2t.net/addgene:41395; RRID:Addgene_41395). pEGFP-c2 from Clontech. GFP-LANA and LANA deletion plasmids were described previously ^40,68^. For generation of MeCP2 T158M mutation, RNA was isolated from LCL 487 (T158M) cell line by Qiagen RNeasy mini kit (cat no # 79656) and cDNA was prepared by maxima H minus first strand cDNA synthesis kit (cat no # K1682). A fragment of MeCP2 containing the T158M mutation was amplified by PCR and cloned in pDEST-hMeCP2-GFP (a gift from Huda Zoghbi, Addgene plasmid # 48078; http://n2t.net/addgene:48078; RRID:Addgene_48078) ^69^and subsequently cloned into pHR-scFv-GCN4-sfGFP-GB1-dWPRE plasmid. MBD domain deleted MeCP2 was created by overlap extension PCR. Oligomerization mutant LANA was created by Q5 site directed mutagenesis kit (cat no# NEB E0554S). For ChIP assays we cloned the dCas9-5xSunTag from pCAG-dCas9-5xPlat2AflD, a gift from Izuho Hatada (Addgene plasmid # 82560; http://n2t.net/addgene:82560; RRID:Addgene_82560) ^70^, and sgRNA for ERBB2 promoter into lentiCRISPRv2 hygro, a gift from Brett Stringer (Addgene plasmid # 98291; http://n2t.net/addgene:98291; RRID:Addgene_98291)^71^. To generate the SunTag only (without dCas9), plasmid pHRdSV40-dCas9-10xGCN4_v4-P2A-BFP was digested by restriction enzymes MluI and BamHI and a linker that also encode for FLAG tag was inserted. All primers can be found in supplementary primer list (**Table S1**).

### Transfection and Immunofluorescence assay

1X10^5^ cells were plated in 12 wells plate. Next day the cells were transfected using PolyJet In Vitro DNA Transfection Reagent (SignaGene Laboratories Cat # SL100688). Each well of a twelve-well plate received 750 ng of plasmid for transfection. The pHRdSV40-dCas9-10xGCN4 v4-P2A-BFP (Addgene plasmid # 60903), scFv-proteinX, and sgRNA telomere plasmids were mixed in a 1:2:4 ratio. In the next day, 4X10^4^ cells were transferred into chamber slide cell culture glass 8 wells (SPL life Science LTD). Next day, the cells were washed once with PBS followed by fixation in 4% paraformaldehyde in PBS pH 7.4 for 10 minutes. After 3 washes with PBS, cells were permeabilized for 10 min with PBS containing 0.25% Triton X-100. Cells were washed 3 times with PBS and blocked with 1% BSA, 22.52 mg/mL glycine in PBST (PBS+ 0.1% Tween 20) for 45 min in 37° C. The slides were incubated for 1 hour with primary antibodies, followed by 3 washes with PBS. Then, the slides were incubated with secondary antibodies for 1 hour and washed again 3 times with PBS and mounted with Vectaschield containing DAPI (Vector Laboratories, H-1500).

### Antibodies

The following antibodies were used for immunofluorescence studies: Rat anti LANA (Advanced Biotechnologies, cat#13-210-100), Mouse anti LANA (Leica, NCL-HHV8-LNA), Mouse anti TRF2 (Merck, 05-521), Rabbit anti MeCP2 (Abcam, ab2828), Rabbit anti mSin3a/SIN3A (Abcam, ab3479), Mouse anti HA (Sigma H9658), Rabbit anti HA (ab9110), Mouse anti ORC2 (MBL Life Science, M055-3), and Rabbit anti HDAC1 (Abcam, ab53091). Goat anti rabbit Alexa 488 (Abcam, ab150077), Goat anti Rat Alexa 488 (Abcam, ab150157), Goat anti Mouse Alexa 594 (Abcam, ab150116) and Goat anti rabbit Alexa 555(ab27039). scFv-MeCP2, scFv-HP1α, and scFv-HDAC1 were detected with Mouse anti HA. scFv-LANA was detected by Mouse anti LANA, except in combination with TRF2 where Rat anti LANA was used. scFv-MeCP2 was detected by Mouse anti HA except for the experiment for the forced dimerization experiment on SunTag in iSLK BAC16 where rabbit anti HA was used. FLAG-LANA mutants (N+C, N, C) were detected with Rabbit anti FLAG (Sigma 7425).

### Confocal imaging

Images were captured with Zeiss LSM780 inverted confocal microscope through a 63X objective with Z stack mode. Then, the middle stacks were selected and merged using Subset mode and Maximum Intensity Projection mode, respectively with Zenn software black edition. Processing and exporting of the resulted images was done with Zenn software blue edition.

### Chromatin immunoprecipitation assay

ChIP assays were performed as described previously ^72^. Immunoprecipitation was performed with protein A/G Dynabeads (ThermoFisherScientific, #100-02D/100-04D) bound with Rat anti-LANA (Advanced Biotechnologies, cat#13-210-100), Rabbit anti-MeCP2 (Abcam, ab2828), or normal IgG (Millipore, PP64B) as a negative control. The immunoprecipitated chromatin was washed five times (with NaCl, lithium chloride and Tris-EDTA buffers), eluted and then purified using QIAquick kit (Qiagen, #28104) according to manufacturer’s instructions.

### LCL infection and maintenance assay

LCLs were infected with KSHV by co-culture ^73^. Briefly, Vero-rKSHV.219 cells (45) were induced with 1.25 mM sodium butyrate and 20 ng/ml 12-*O*-tetradecanoylphorbol-13-acetate (TPA) and after 48 h were cocultured with LCL cells (1:1 cell ratio) in RPMI 1640 medium for an additional 48 h. Medium containing the LCLs infected cells was collected and transferred to a new flask for an additional 48 h. Transferred again to a new flask, and at days 2, 7, 14, 24, cells were taken for FACS analysis to detect the number of GFP positive cells. At the same time points, genomic DNA was isolated with DNeasy blood and tissue kit (cat no # 69506) as per manufacturer instructions. Real time PCR was performed on genomic DNA using Fast SYBR green master mix (Applied Biosystems), and results were analyzed with a CFX96 Touch real-time PCR detection system (Bio-Rad). GAPDH was used as an internal control.

### shRNA transduction and colony assay

shRNA for MeCP2 and shRNA control were ordered from horizon discovery and named as shMeCP2#1(V3LHS_410280), shMeCP2#3(V3LHS_378909), and shControl (Non-silencing Verified Negative Control, RHS4346). The shRNA lentiviruses were prepared in HEK 293T cells as per manufacturer’s protocol. Briefly, 10 cm dish of 293T cells were co-transfected with 2 μg of respective shRNA plasmid, 2 μg psPAX2, and 1μg pMD.2G packaging plasmids. Day after transfection, old media was replaced with new 5ml DMEM medium. 48h after transfection, supernatant containing lentivirus was collected and centrifuged at 1200 rpm for 5 mints at 4°C and filtered through 0.45um filter. The filtered lentiviral supernatant was then added to iSLK BAC16 cells in the presence of 4ug/ml polyberene and incubated for 4h. Then lentivirus containing media was removed and fresh DMEM medium containing 1200ug/ml hygromycin was added on iSLK BAC16 cells. 48h after transduction, iSLK BAC16 cells were trypsinized, counted and around 3000 cells were plated in 10 cm dish in the presence of 1200ug/ml hygromycin for colony formation assay. Hygromycin was replenished twice a week in fresh media until individual clones grow out after ∼10days. Cell colonies were stained with gentian violet (MP biomedicals cat no # 0210177525) and counted with ImageJ software.

### Statistics

Statistical significance of the results were calculated in Prism 9 (GraphPad). Pearson’s correlation coefficient was determined by ImageJ (JACoP Plugin) ^74^.

## Data availability

N.A., as we did not generate large data sets.

## Acknowledgments

We would like to thank J Ron Vale, Stanley Qi, David Root, Izuho Hatada, Brett Stringer, and S. Diane Hayward for kindly providing plasmids, and Don Ganem, Rolf Renne, Jeffrey Vieira, Oren Kobiler, Richard F. Ambinder, Bert Vogelstein, Jae U. Jung, Ronit Sarid and Uta Francke for cell lines. We also would like to thank S. Diane Hayward for critically reading the manuscript, and Yosef Shaul for fruitful discussions.

