## supplementary figures and table for "Unidirectional protein recruitment is essential for viral latency"

Ido Lavi, et al.

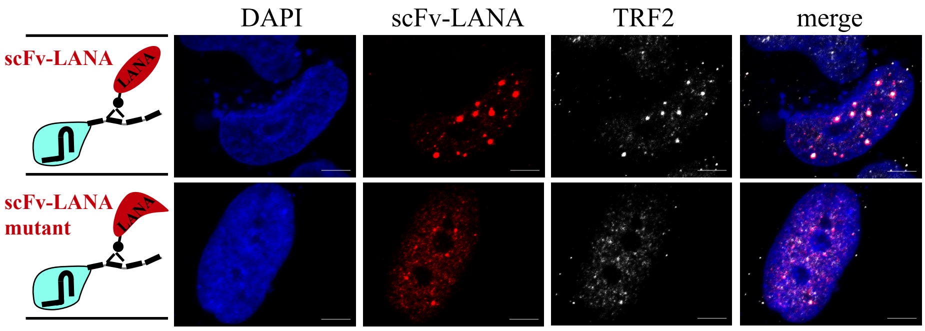

**FIG S1. LANA-telomere dots co-localize with the telomeric protein TRF2.** SLK cells were transfected with dCas9-SunTag, sgTelomere, and scFv-LANA or scFv-LANA oligomerization mutant as illustrated on the left of the images. Immunofluorescence assay was performed to detect LANA (red) and TRF2 (white), and the nucleus was stained with DAPI. Images are representatives of at least two independent experiments. Scale bar = 5 µm.

**Fig. S1**

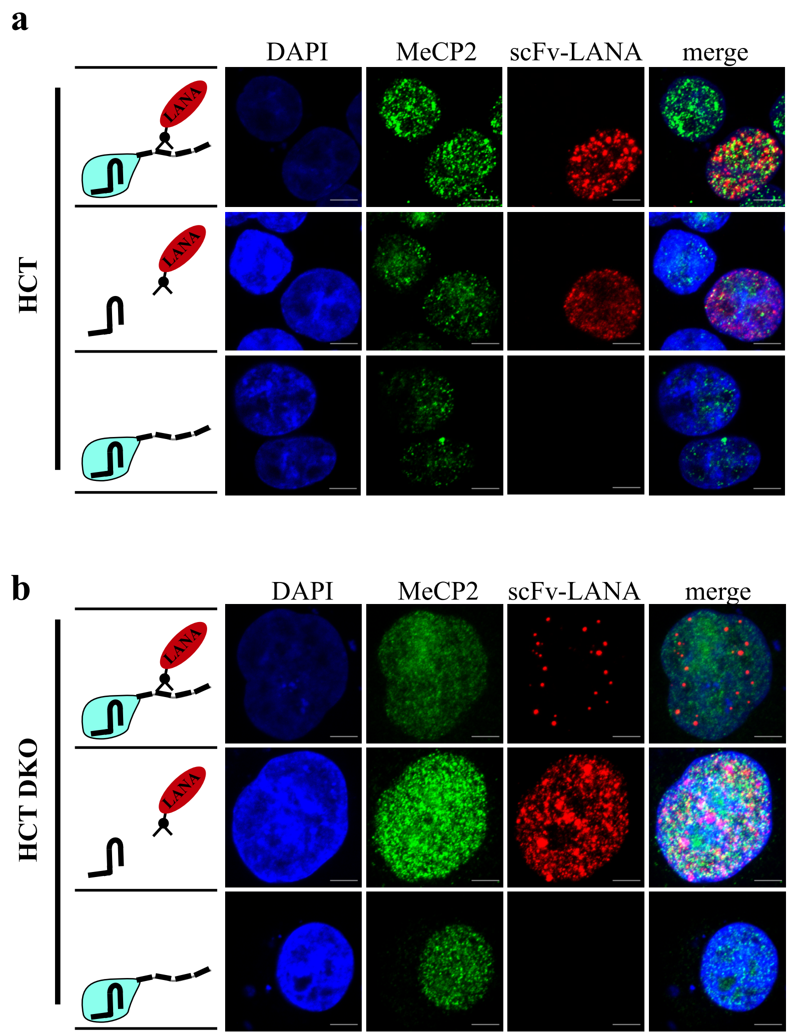

**FIG S2. LANA cannot recruit MeCP2 even in HCT-DKO cells.** HCT116 (A) and DKO (DNMT3B-/- and DNMT1 -/-) HCT116 cells (B) were transfected with dCas9-SunTag, scFv-LANA, and sgTelomere as illustrated on the left of the images. Immunofluorescence assay was performed to detect MeCP2 (green) and LANA (red). The nucleus was stained with DAPI. Images are representatives of at least two independent experiments. Scale bar = 5 μm.

**Fig. S2**

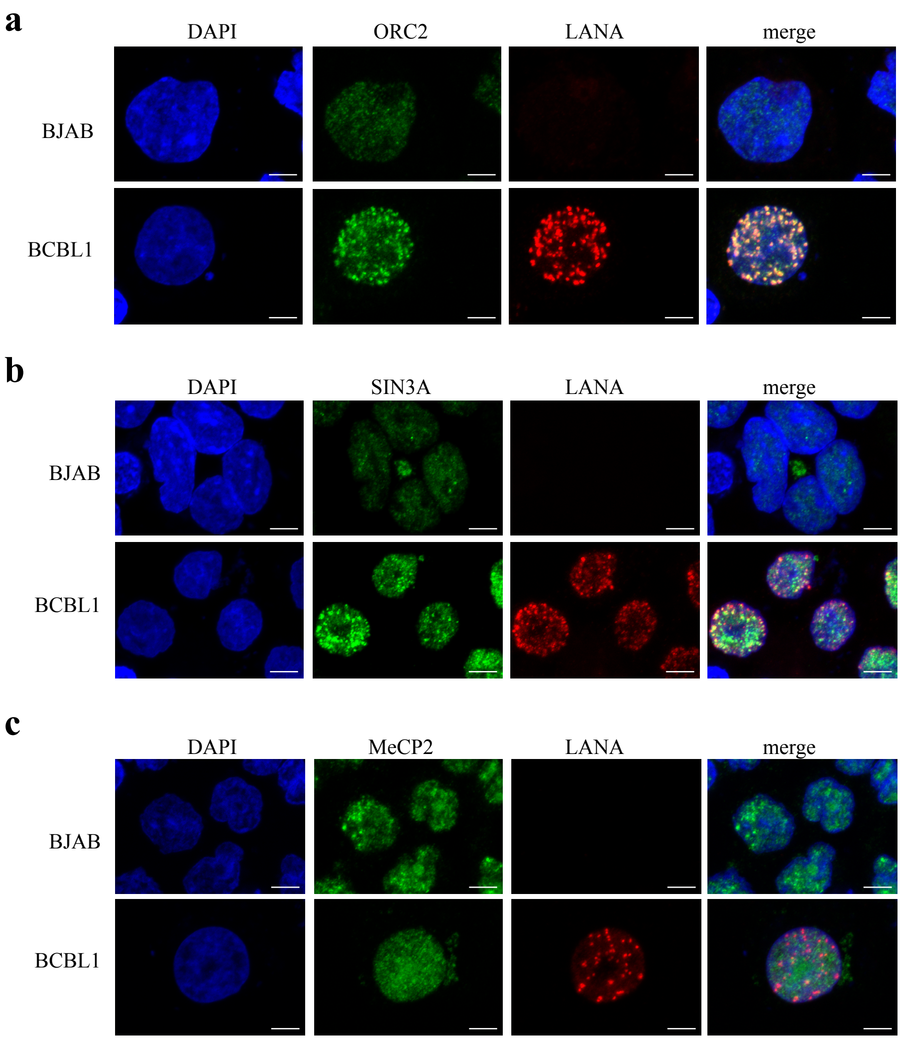

**FIG S3. ORC2 and SIN3A but not MeCP2 co-localize with LANA dots in KSHV infected cells.** Immunofluorescence assays were performed to detect ORC2 (**A**) SIN3A (**B**) or MeCP2 (**C**) cellular localization in KSHV-negative (BJAB) and KSHV-positive (BCBL1) lymphoma cell lines. The nucleus was stained with DAPI. Images are representatives of at least two independent experiments. Scale bar = 5 µm.

**Fig. S3**

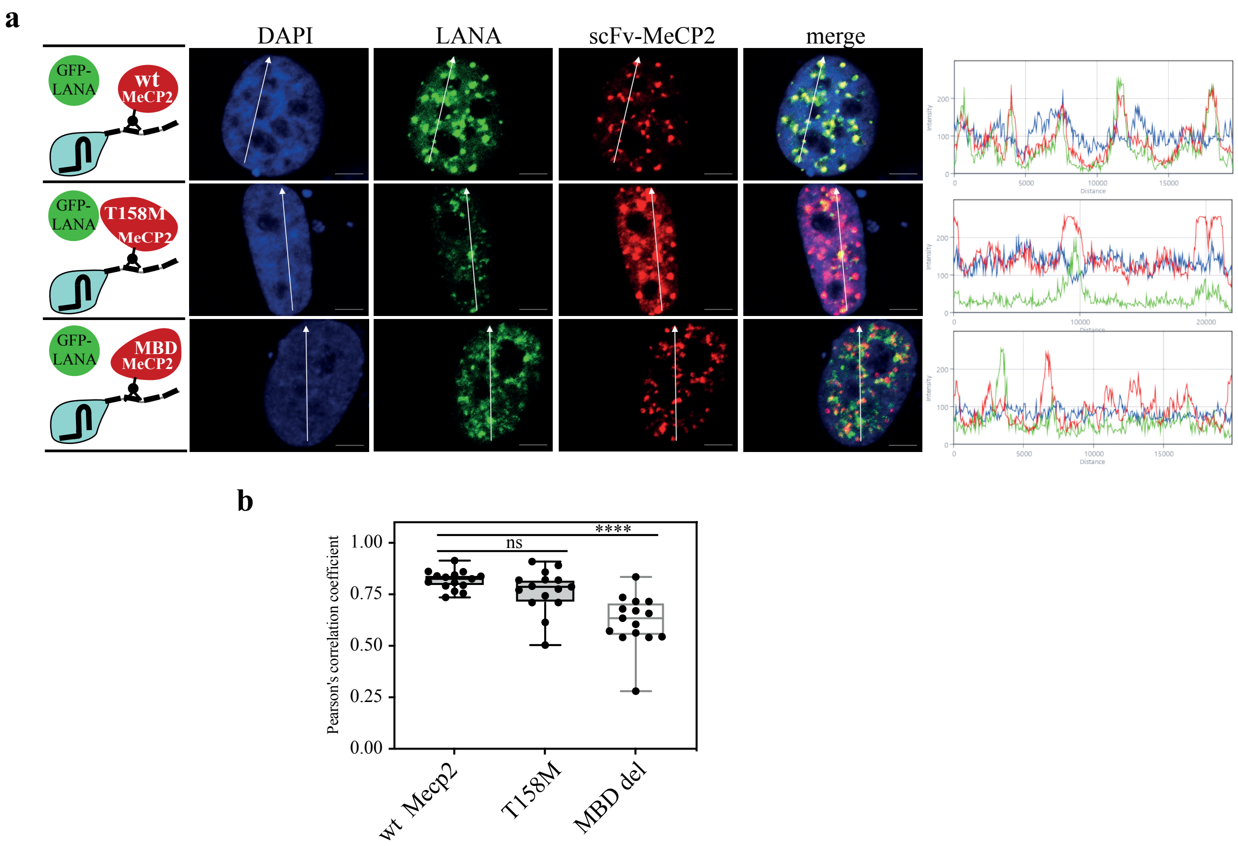

**FIG S4. MBD deletion abrogates the ability of MeCP2 to recruit LANA. (A)** SLK cells were transfected with dCas9-SunTag, scFv-MeCP2, GFP-LANA and sgTelomere. Different scFv-MeCP2 constructs were transfected to express wild-type (wt), T158M mutant or a MBD deletion mutant of MeCP2, as illustrated on the left. Immunofluorescence assay was performed to detect scFv-MeCP2, or fluorescently labelled GFP-LANA. The nucleus was stained with DAPI. Scale bar = 5 µm. The plots of the red, green and blue pixel intensities along the white arrow (in the left panels) are presented on the right. Images are representatives of at least three independent experiments. (**B**) Pearson's correlation coefficient was determined for 15 different cells in each treatment, and presented as box and whiskers (min to max). One-tailed *t* tests were performed (*, P ≤ 0.05; **, P ≤ 0.01; ***; P ≤ 0.001).

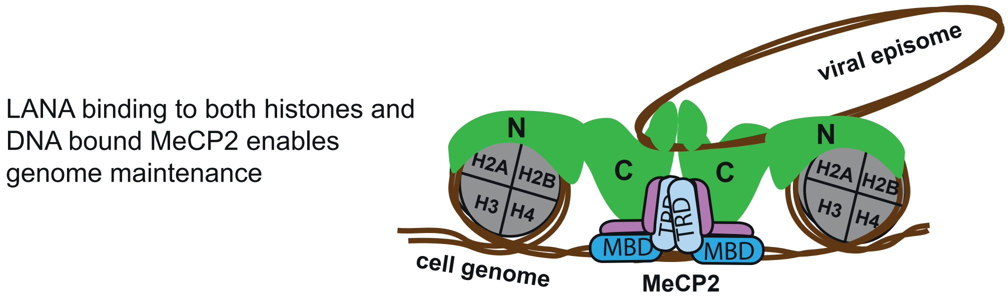

**FIG S5. Schematic presentation of tethering by LANA.** LANA tethers the viral episomal genomes to cellular chromosomes by binding to histones via its N-terminal domain and to DNA bound MeCP2 via its C-terminal domain. The unidirectional recruitment of LANA by MeCP2 is essential for viral genome maintenance during latency.

**Fig. S3-S4**

**Table S1. Primers used in this study.**

| | |
|---|---|
| **Primer used for KSHV episome maintenance** | |
| LANA F | GAAGTTGTAGGAAACGAAACAGGT |
| LANA R | ACACTGTGGGACTTCCAGGTATAG |
| GAPDH F | CCTCAAGCAC CACTTTGTCA |
| GAPDH R | CCCTGTTGCT GTAGCCAAAT |
| **Primer used for ChIP** | |
| ERBB2a Chip F | AGTCACCAGCCTCTGCATTTA |
| ERBB2A Chip R | CCAGCTTCACTTTCTCCCTCT |
| **Primer used for cloning** | |
| LANA mut F | gacgcgcTTTGGGAAAGGATGGAAGAC |
| LANA mut R | tataagcGATGTGTTGTGGCCTAGC |
| HDAC1 F | GATCCgATGGCGCAGACGCAGG |
| HDAC1 R | GCTCAGGCCAACTTGACCTCCT |
| HP1aF | GAGGGATCCGATGGGAAAGAAAACCAAGCGGAC |
| HP!a R | TCTGCGGCCGCTTAGCTCTTTGCTGT |
| MeCP2F | GAGGGATCCGATGGTAGCTGGGATGTT |
| MeCP2R | AGAGCGGCCGCCTAGCTAACTCTCTCG |
| MECP2 (T158M) F | gacacggaagcttaagcaaaggaaa |
| MECP2 (T158M) R | ttgggaatggcctgaggg |
| MBDdelFRAG1F | gcttgGATCCgatggtagctg |
| MBDdelFRAG1R | aggggctcccagaagcttccggcacagccg |
| MBDdelFRAG2F | gaagcttctgggagcccctcccgg |
| MBDdelFRAG2R | agtcgcGGCCGCct |
| sgRNA telomere | TTGGGTTAGGGTTAGGGTTAGGGTTAGTTTTAGAGCTAGAAATA GCAAGTTAAAATAAGGCTAGTCCGTTATCAACTTGAAAAAGTGG CACCGAGTCGGTGCTTTTTTC |

**Table S1. Primers used in this study.**

| | |
|---|---|
| sgRNA ERBB2 F | CACCGGTTGCCACTCCCAGACTTGT |
| sgRNA ERBB2R | AAACACAAGTCTGGGAGTGGCAACC |
| MluI FLAG BamHI S | cgcgacGATTACAAGGACGACGATGACAAGa |
| MluI FLAG BamHI AS | gatctCTTGTCATCGTCGTCCTTGTAATCgt |
